# Mutalyzer 2: Next Generation HGVS Nomenclature Checker

**DOI:** 10.1101/2020.06.24.168583

**Authors:** Mihai Lefter, Jonathan K. Vis, Martijn Vermaat, Johan T. den Dunnen, Peter E.M. Taschner, Jeroen F.J. Laros

## Abstract

Unambiguous variant descriptions are of utmost importance in clinical genetic diagnostics, scientific literature, and genetic databases. The Human Genome Variation Society (HGVS) publishes a comprehensive set of guidelines on how variants should be correctly and unambiguously described. We present the implementation of the Mutalyzer 2 tool suite, designed to automatically apply the HGVS guidelines so users do not have to deal with the HGVS intricacies explicitly to check and correct their variant descriptions. Mutalyzer is profusely used by the community, having processed over 133 million descriptions since its launch. Over a five year period, Mutalyzer reported a correct input in approximately 50% of cases. In 41% of the cases either a syntactic or semantic error was identified and for approximately 7% of cases, Mutalyzer was able to automatically correct the description.

## 1 Introduction

The Human Genome Variation Society (HGVS) publishes^1^ nomenclature guidelines [1, 2, 7, 8] for the unambiguous description of genetic variants in order to prevent undesired errors in clinical diagnostics and to enable sharing and comparison of variants across different institutes. Since their initial introduction, the HGVS guidelines are continuously extended and adapted to accommodate the evolution of the domain. As a result, it has become increasingly complex for users, i.e., researchers, database curators, and manuscript reviewers, to check whether variant descriptions comply with the HGVS guidelines.

A method that automatically deals with the HGVS intricacies and outputs correct unambiguous variant descriptions is of high necessity for consistent variant dissemination. In this context, the Mutalyzer tool suite was created to assist geneticists in applying the HGVS guidelines in databases (e.g., [9, 5, 18]) and literature^2^ by providing the means for automatic checking and correction of HGVS variant descriptions.

In this paper we present the second iteration of the Mutalyzer suite. While bearing the same name as its initial version [20], Mutalyzer 2 is a complete new implementation of this idea. Its core functionality is provided by the *Name Checker* tool which provides checking and disambiguation of variant descriptions. The Name Checker is able to process variant descriptions for any organism, as long as the corresponding reference sequences can be retrieved from supported sources. In addition, it offers features such as protein effect prediction in the form of amino acid changes relative to the protein reference sequence. Several other related tools are included in the suite, e.g., the *Position Converter*, which converts descriptions between chromosomal and transcript references and the *Description Extractor*, which computes a description when provided with two sequences.

## 2 Problem Description

A *variant* in the context of molecular sequence analysis is the difference between two sequences, i.e., strings over a molecular alphabet. One of them is chosen to be the *reference sequence*, the other one is named the *observed sequence*. Given a reference sequence and a variant, the observed sequence can be reconstructed. Variants can be expressed in different *description languages* [3, 6, 8, 11], among which various inconsistencies exist. In the following we consider the HGVS nomenclature [8] as a description language for variants.

The representation of a variant in the HGVS description language [13] is a list of *elementary variants*, i.e., transformations that together, not independently, describe the difference between the reference and the observed sequence. Within the HGVS description language, the same variant can be described in different ways. We define the set of *interpretable* descriptions of a variant as all equivalent ways of describing the same variant.

The HGVS nomenclature offers guidelines to select a *canonical* variant description from the set of interpretable descriptions, i.e., to *disambiguate* a description. However, a deterministic algorithm for doing so is not given, instead, numerous examples are given from which an attempt of creating such an algorithm can be made. The challenge is to create an algorithm that accepts all interpretable HGVS variant descriptions and finds the canonical one.

## 3 HGVS Variant Descriptions

An HGVS variant description is composed of three parts: a *reference sequence identifier*, a *positioning system* and a list of elementary variants. Each elementary variant consists of the positions of the reference sequence that are affected, the operation to be performed and, optionally, what is to be newly inserted.

Consider the following description of an allele written in the HGVS language, which is interpretable but not canonical.

~~~
NG_012337.1:g.[7G>T;14del].
~~~

Here, the reference sequence identifier is `NG_012337.1`, the positioning system is `g.` (linear genomic) and the elementary variants are `7G>T` and `14del`. The first elementary variant indicates that at position 7 instead of a `G`, a `T` was observed, while the second indicates that the nucleotide present at position 14 in the reference sequence is missing from the observed sequence. A visual representation is given in Figure 1(a).

**Figure 1:**
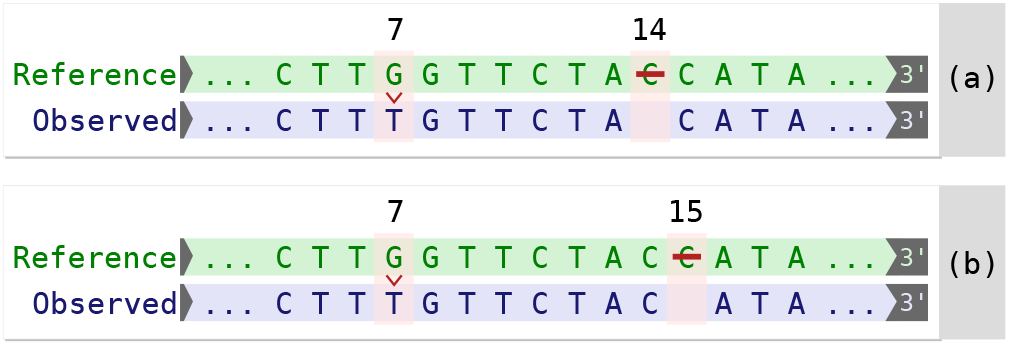
Elementary variants of descriptions `NG_012337.1:g.[7G>T;14del]` (a) and `NG_012337.1:g.[7G>T;15del]` (b) applied to a reference sequence in order to obtain the same observed sequence.

Ambiguity arises if multiple descriptions lead to the same observed sequence. In the example given above, a C occurs at positions 14 and 15, either of which can be deleted in order to be left with one C in the observed sequence. Consequently, the description [7G>T;15del] leads to the same observed sequence, as depicted in Figure 1(b). According to the HGVS recommendations the latter description is the preferred description since it respects the 3’ rule [8]. Note that in the Variant Call Format [6], a left shift with respect to the genome should be performed [21], there 14del is preferred over 15del.

## 4 Approach

Given an interpretable description, the Mutalyzer *Name Checker* finds the canonical description using the following general approach. First, a syntactic check is performed, which allows for the interpretation of the description and the retrieval of the reference sequence. Next, a semantic check is done to establish whether the description makes sense in the context of the reference sequence. Finally, the description is disambiguated.

### 4.1 Preliminaries

The formalisation of the HGVS syntax [13] has made it possible to implement a context free grammar parser, which recognises a syntactically valid description and generates a *parse tree*. In this manner, all parts of the description can be accessed programmatically.

If the variant description is syntactically correct, i.e., it can be parsed, its validity in the context of the reference sequence is verified. The reference sequence, identified by the accession and version numbers, is retrieved from a reference sequence repository. Currently, the supported sources are the National Center for Biotechnology Information (NCBI) (for GenBank files [15]) and the European Bioinformatics Institute EMBL-EBI (for Locus Reference Genomic (LRG) files [14]). Reference sequence files contain a reference sequence and feature annotations, e.g., locations of transcripts, coding sequences (CDS) and exons.

To work with different types of reference sequence files, an abstraction layer is used. For every reference sequence file type, a dedicated parser provides a uniform *reference model* that can be used for the semantic check. Because the annotation of reference sequence files is not always complete, an annotation enrichment procedure is used to standardise this information. Annotation enrichment consists mainly of the linking of transcripts to its CDS and protein. In many cases, (especially in GenBank files) there is no direct link between a transcript and its CDS, making it impossible to reconstruct the layout of the transcript and thereby the biological effect of a variant. The enrichment procedure will try to deduct these links by other means (e.g., by additional queries to the NCBI databases).

The HGVS nomenclature supports different positioning systems [8]. In contrast to most positioning systems used in mathematics and computer science, the number zero is not included in any of the HGVS positioning systems. As a result, in the `c.` system, which can be regarded as the most complex one, positions `-1` and `1` are adjacent, introducing a discontinuity. Additionally, the first position after the CDS is denoted as `*1`, introducing another discontinuity. Finally, the use of *offsets* for intronic positions introduce yet another type of discontinuity, e.g., position `c.12` can be adjacent to `c.12+1` while it is not adjacent to `c.13.` For transcripts that reside on the reverse complement strand, the direction of their positioning system is opposite to that of the genomic one, e.g., if `c.1` is equivalent to `g.10`, then `c.2` is equivalent to `g.9.` These peculiarities make performing arithmetic operations in the HGVS positioning systems tedious and error prone. Mutalyzer uses an internal zero-based half-open coordinate system and implements the conversion from HGVS `c., g.`, and `n.` positioning systems to this coordinate system and vice versa in order to perform arithmetic operations. Note that the HGVS genomic circular `o.` positioning system^3^ is currently not supported and that the special mitochondrial DNA `m.` is treated in a similar manner as `g.`.

### 4.2 Semantic Checking

With the parsed description and the information from the reference model, a semantic check can be performed. This is done in multiple steps, described below.

If a transcript ID is provided and its annotation is present in the reference sequence file, the corresponding exons and CDS positions are retrieved and used to convert the positions of the elementary variants to the internal coordinate system. This allows for the verification of the given positions from different perspectives. First, all positions should be within the sequence boundaries. Second, for ranges, the end position should be greater than the start position. Third, for insertions, the start and end positions should be consecutive. Fourth, offsets for intronic positions should start on exon boundaries and should have the correct orientation following the annotation in the reference model.

To better assist users, Mutalyzer still supports several deprecated HGVS constructs. Old intronic position descriptions, such as `c.IVS4+1`, which referred to the first nucleotide of intron 4, are converted to the correct format. In addition, if a range length is provided, its length is checked for equality to the length determined from the positions, e.g., `3_9del7.` When deleted sequences are specified in a description, such as `10_12delAAT`, Mutalyzer checks whether the reference sequence indeed contains the sequence `AAT` in the indicated range. No checks are performed on specifications of inverted sequences.

### 4.3 Disambiguation

After the syntactic and semantic checks, disambiguation is performed. First, a check is performed for the `delins` and `inv` elementary variants in order to determine whether the minimal description is used.

For example, for the reference sequence `AACGTAA`, the deletion-insertion `3_4delinsTT` results in the observed sequence `AATTTAA`. The same result is obtained when the variant `2_6delinsATTTA` is applied. The latter description can be minimised by removing the *longest common prefix* and the *longest common suffix* of the deleted and the inserted sequence.

For an inversion, a prefix of the inversion can be equal to the reverse complement of its suffix, i.e., a *partial palindrome*. The description of an inversion is minimised in a similar way as described above.

After the minimisation step, a simplification scheme is used to check whether the elementary variant type used is the simplest one possible. A `delins` can insert a sequence that is a prefix or a suffix of the deleted sequence. In this case, the `delins` should be simplified to a `del`. Likewise, if the deleted sequence is a prefix or suffix of the inserted sequence, the `delins` is actually an `ins`. For example, given the reference sequence `AACGTAA`, the deletion-insertion `2_5delinsGT` is simplified to `2_3del` since the inserted part, `GT` is a suffix of the deleted part, `ACGT`. For the same reference sequence, `3delinsT` is simplified to the substitution `3C>T`.

Similar checks are performed for other cases, the elementary variant types that can possibly be written as a simpler type are shown in Table 1.

**Table 1:**
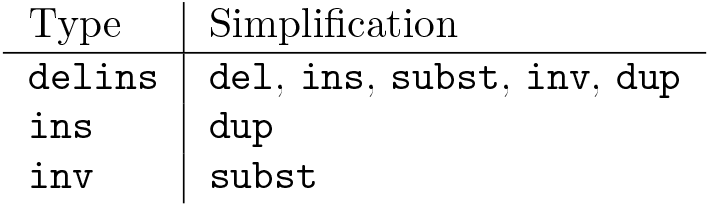
Simplification of elementary variant types.

Finally, a deletion, insertion or duplication is shifted to the most 3’ position possible. An algorithm that considers all circular permutations of the deleted or inserted sequence is used in order to find this position. If for example, the sequence `CGTC` is inserted in the reference sequence `AACGTAA`, the algorithm will correct the description `2_3insCGTC` to `5_6insCCGT`.

Note that this method is applied to both the forward as well as the reverse strand, so if a gene resides on the reverse strand, the position will be shifted in the opposite direction to that of the genomic one. Furthermore, if an insertion or a deletion is described on a transcript, the position will not be shifted over a splice site.

## 5 Related Problems

In addition to variant description disambiguation, Mutalyzer tackles some related problems in order to aid the user with the interpretation and validation of variant descriptions.

### 5.1 Effect Prediction

Apart from the effect a variant has on a transcript, the impact on the potential protein is also highly relevant. Therefore, the Name Checker runs a series of effect preditions after a successful, i.e., error free, disambiguation step.

For all the annotated transcripts in the reference sequence, the corrected variant description is shown together with a list of protein descriptions. Each of the variant descriptions can be selected for a more detailed analysis. In the detailed analysis, the reference protein and the variant protein are visualised with the area of change highlighted. Additionally, for the selected transcript, a list of exon start and end positions is given, as well as the CDS start and end positions.

For all elementary variants, effects on restriction sites are calculated. A table is generated that contains a list of removed restriction sites and a list of added restriction sites for every elementary variant.

Effect prediction becomes more complex when variants are near or overlapping splice sites. Barring some specific cases, Mutalyzer takes a safe approach by issuing a warning and omitting prediction of a translated protein if a splice site is hit, e.g., `NG_012772.3(NM_000059.3):c.508_516+9del`. However, a deletion partly covering two exons, thereby spanning an intron, although affecting two splice sites, is considered to be a case where protein prediction can still be valuable. For example, for `NG_012772.3(NM_000059.3):c.508_525del`, the deletion is interpreted as forming a fusion exon. Likewise for the deletion `NG_012772.3(NM_000059.3):c.317-10_631+14del`, covering four complete exons, Mutalyzer removes the exons from the CDS to give a meaningful protein prediction. In practice, large deletions are often described by using fuzzy intronic offsets (e.g., `NG_012772.3(NM_000059.3):c.317-?_631+?del`). Internally and for the conversion to `g`. descriptions, Mutalyzer interprets these offsets as being in the center of the intron.

### 5.2 Support for contigs and chromosomes

Large GenBank files, for instance whole chromosomes or contigs, cannot be parsed in a reasonable amount of time, which makes on-line parsing during the page load of an interactive web page impractical. For this reason, the maximum accepted file size for the Name Checker is set to 10 MB. To be able to work with larger reference files, the *Reference File Loader* can be used to generate slices of large reference files which are subsequently uploaded to Mutalyzer in order to be used in variant descriptions. These references are provided with an internal accession number starting with `UD_`.

A slice can be made directly by supplying the name or accession number of a chromosome or contig, the slice start and end positions, and the orientation. Alternatively, a gene name in combination with the name of an organism and the sizes of the flanking regions can be used to select the slice automatically from the latest genome build of the organism.

Since the generated reference identifiers are only valid within a particular Mutalyzer instance, using these identifiers for dissemination is not recommended. This is why Mutalyzer additionally provides the genomic variant description with respect to the origin of the slice. This workaround has been superseded by full support for chromosomal references described in Section 5.2.2.

#### 5.2.1 Transcript and chromosome mappings

The *Position Converter* converts a description using a RefSeq transcript to one using a chromosomal reference sequence and vice versa. For this purpose it uses mapping information that is retrieved from the NCBI. Currently human genome builds NCBI36, GRCh37 and GRCh38, as well as mouse (GRCm38) and dog (canFam3) are supported.

The Position Converter can be used to quickly convert variants found by a high throughput screening technique like Next Generation Sequencing to transcript oriented HGVS descriptions. Another use of this interface is to convert (lift over) a description from one transcript to another, or to transcripts of other (overlapping) genes. Finally, by using transcripts that are mapped to multiple genome builds, it is possible to convert a chromosomal description from one build to another. Potentially, descriptions can be lifted over to other species, provided cross-species transcript annotation is available.

Note that the Position Converter does not perform any semantic checks or disambiguations, nor does it take any differences between reference sequence content into account (for a discussion on the underlying problem, see Section 7.1). This is why the output of the position converter should always be checked with the Name Checker before dissemination. In Section 5.2.2, we provide an alternative to this procedure.

#### 5.2.2 Support for chromosomal references

Support for full chromosomal references has been lacking until recently because of the time consuming nature of reference sequence file parsing. Since version 2.0.27 however, Mutalyzer is able to preprocess reference sequences of human genome builds GRCh37 and GRCh38, the results of which are stored in a database for quick retrieval. This allows for the on-line handling of HGVS descriptions for full chromosomal references of the aforementioned genome builds.

With this added functionality, users do not have to rely on the Reference File Loader or the Position Converter to analyse variants described on chromosomes.

### 5.3 Generating Descriptions

While checking the validity of variant descriptions is important, methods aiding the user in the generation of such descriptions is at least as valuable, especially for those who are new to the field. Mutalyzer addresses this topic from two different perspectives.

#### 5.3.1 Name Generator

The *Name Generator* is an interactive interface for the construction of HGVS variants, tailored to people that are less familiar with the HGVS nomenclature. After choosing a reference sequence, variants can be added one by one in an easy to use point and click interface. The HGVS variant description is build incrementally from these variants and can be checked with the Name Checker afterwards.

#### 5.3.2 Description Extractor

A recent addition to the Mutalyzer tool suite is the HGVS variant *Description Extractor* [19]. This tool automatically generates HGVS variant descriptions given a reference sequence and an observed sequence. As a deterministic algorithm for the generation of variant descriptions, this method will also be applied in the disambiguation of complex variant descriptions when using the Name Checker. Currently, the description extractor runs as an experimental service in this context.

## 6 Usage

In this section, we present the interfaces that provide access to the tools in the Mutalyzer suite. Furthermore, we provide some quantitative insights with respect to the usage of the tools and their interfaces for the LUMC hosted Mutalyzer instance. Note that other instances exist, since Mutalyzer is open source and permissively licensed, hence it can be downloaded and installed locally.

### 6.1 Interfaces

The Mutalyzer *website* interface^4^ provides interactive access to all the tools in the suite. To facilitate automated use of the Mutalyzer tools, two other interface are available.

#### Batch Jobs

The *Batch Checker* is an interface used for processing batches of data in a non-interactive way. It is available for the Name Checker, the Syntax Checker and the Position Converter.

The Batch Checker accepts three types of input formats: CSV files (the delimiters are detected automatically), Microsoft Excel files and OpenOffice ODS files. Each row consists of a variable number of fields, where every field contains a single variant description. For backwards compatibility, the format used by Mutalyzer 1.0.3 is also accepted. The output of a Mutalyzer batch run is a tab delimited CSV file. Note that empty lines are removed from the batch output file. When a submitted batch job is finished, the user receives a download link via e-mail.

#### Web Services

A large number of web services are available to facilitate developers that want to use the Mutalyzer functionality. Currently, two major protocols are supported: SOAP [4] and JSON-RPC [12] over HTTPS. Documentation of the APIs as well as example client scripts in various languages are available on the website^5^.

### 6.2 Statistics

Simple usage statistics are shown in Figure 2, with the number of variant descriptions given as input (or as output in case of the Description Extractor) on the y-axis. From this graph, it can be seen that the Position Converter is the most frequently used tool, irrespective of the interface through which it was accessed, followed by the Name Checker. Also note that the automated interfaces are used more than the interactive ones, with the batch jobs as the most popular interface, followed by the web services and finally the website.

**Figure 2:**
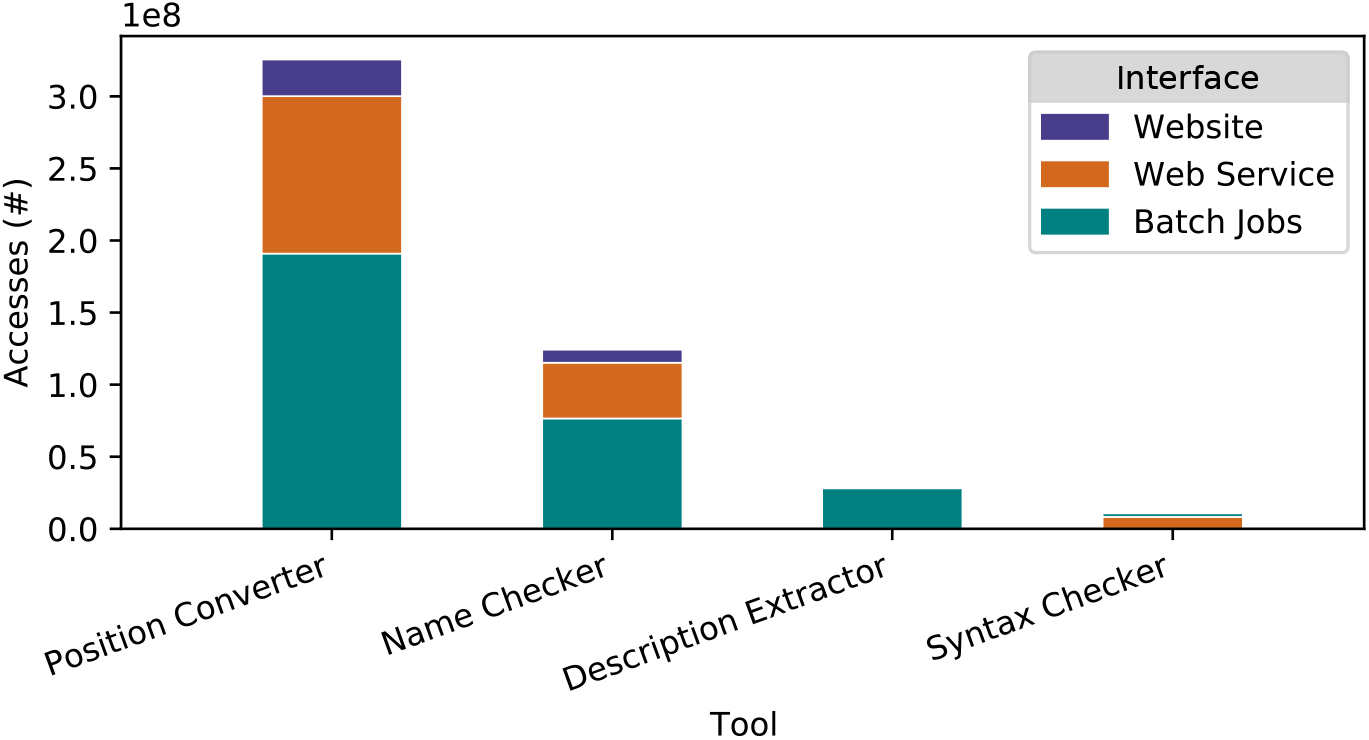
Mutalyzer usage per tool and interface, extracted from https://mutalyzer.nl in December 2019.

In the remainder of this section, we present usage information extracted from the Mutalyzer access logs recorded between September 2014 and October 2019. We focus on the Name Checker and Position Converter.

#### 6.2.1 Name Checker

The Name Checker processed approximately 87 million descriptions in total, of which 20% (17 million) were unique. From the remaining 80%, the most frequent descriptions are invalid or syntactically incorrect, with various forms of notation indicating that no variant is present accounting for approximately 11% of the total.

In total, approximately 26 million unique descriptions were processed by Mutalyzer (all tools). These descriptions were submitted to the Name Checker of a local Mutalyzer 2.0.32 instance to obtain detailed information on the assessment made.

An overview of how the Name Checker assessed the variant descriptions is presented in Figure 3. In 50.4% of the cases, no error or warning messages were reported, signifying a correct input. In approximately 1.2% of the cases, the input was correct, but the Name Checker warned for possible issues such as variants that occur near a splice site or cover the translation start site.

**Figure 3:**
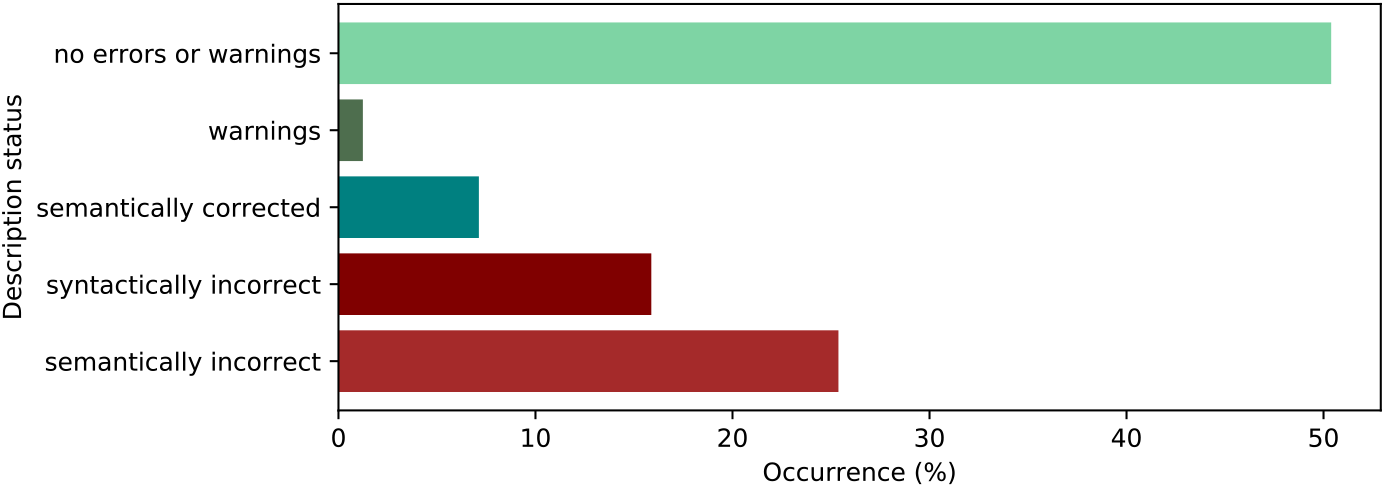
Assessments made by the Name Checker for all submitted descriptions.

The Name Checker had to correct and canonise variant description in approximately 7.1% of the cases, reporting an appropriate warning message related to the operations performed to correct the description (see Section 4). In 36% of the cases the HGVS 3’ rule was applied to disambiguate the input description. In 35% of the cases an accession was given without a version number and the Name Checker had to retrieve the most recent version. Finally, the variant type was updated according to the HGVS prioritisation rules in 29% of the cases. Out of the latter, in more than 65% of the cases an ins was simplified to a dup.

Approximately 15.8% of the descriptions, of which 95% were submitted via the web service interface, did not pass the syntactic check (see Section 4). About 85% of the descriptions submitted via the website were syntactically correct, while descriptions submitted in batch jobs were syntactically correct in only 80% of the cases.

Finally, 25.4% of the descriptions did not pass the semantic analysis of the Name Checker. The most common error messages are shown in Figure 4. The error ‘ENOINTRON’ is the most frequent one (36%), which occurs when an intronic position is used with a reference that does not contain the intronic sequence (see Section 7.1 for a discussion on this topic). Other frequent errors were ‘EREF’ (15%), which is raised whenever a sequence in the description sequence does not correspond to what is present in the reference sequence and ‘EOFFSETFROMBOUNDARY’ (10%), which indicates that an intronic position did not start at an exon boundary.

**Figure 4:**
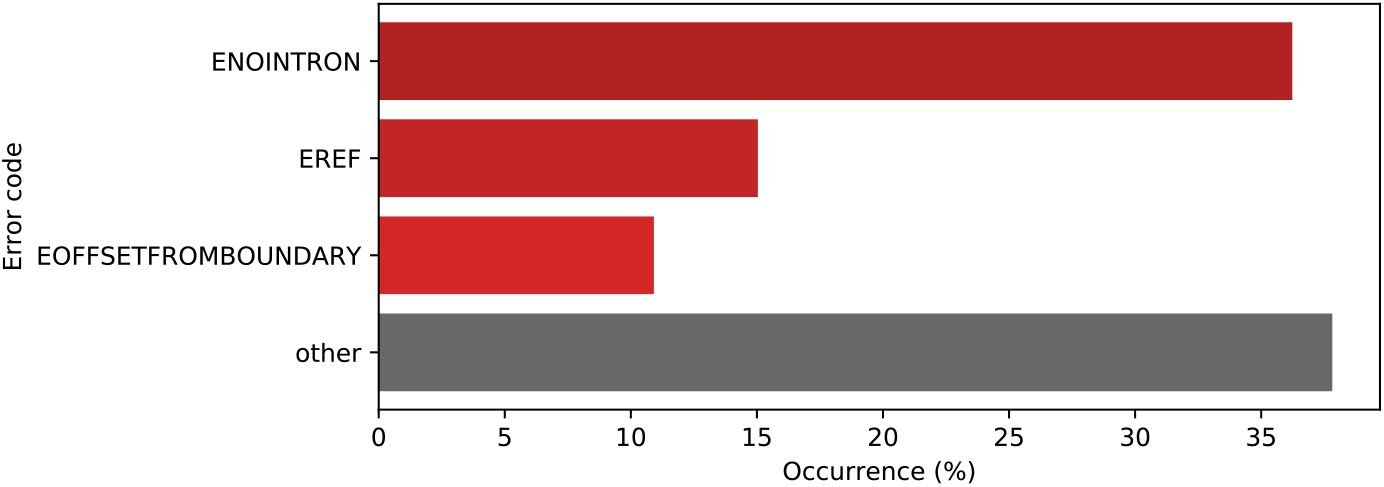
Common error codes returned by the Name Checker.

The most common warnings are shown in Figure 5. ‘WSPLICE_OTHER’ is the most frequent one (59%), occurring whenever a variant is near any of the splice sites of an annotated transcript. Other frequent warnings were ‘WNOM-RNA_OTHER’ (7.7%), which is emitted whenever the transcript model could not be directly retrieved from the reference annotations and it had to be reconstructed from the CDS, and ‘WSPLICE’ (7.6%), which indicates that an elementary operation affects a splice site.

**Figure 5:**
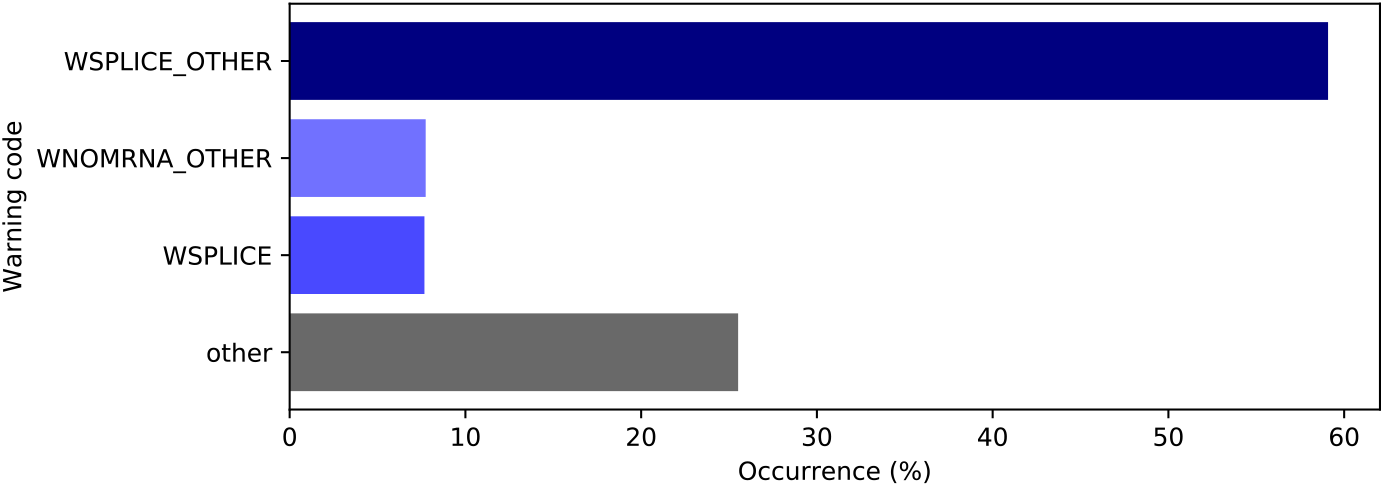
Common warning codes issued by the Name Checker.

The reference sequence types used in syntactically correct descriptions are presented in Figure 6. mRNA reference sequences (prefixed by `NM_` and `XM_`), are used almost twice as much as genomic reference sequences (prefixed by `UD_, NG_, LRG_`, and `NC_`). Since support for full chromosomal (`NC_`) reference sequences was added only recently, `UD_` prefixed (see Section 5.2) are the most used genomic reference sequences. With respect to positioning systems, `c.` was used in more than 90% of the cases.

**Figure 6:**
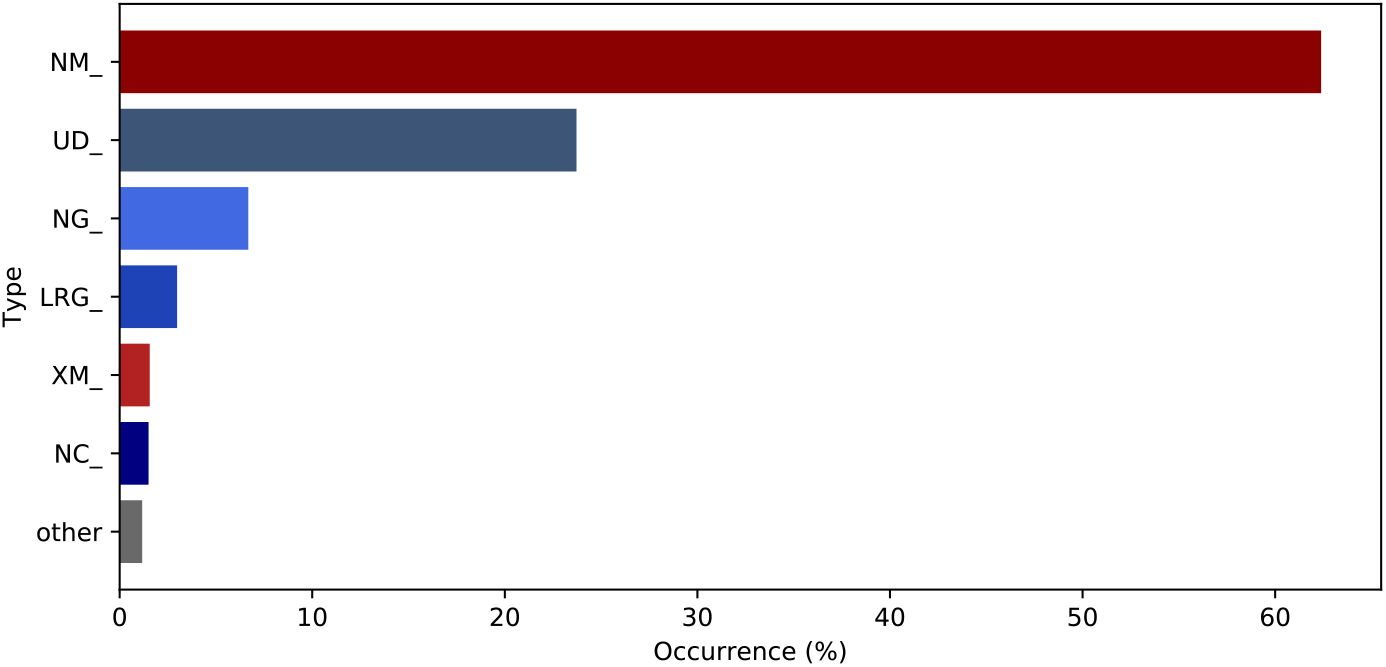
Usage of reference sequence types.

Of all variants encountered, 78% were substitutions, 12% were deletions, 5% were insertions. Duplications and deletion-insertions accounted for 2% each and in 1% of the cases, no variant was provided.

#### 6.2.2 Position Converter

The Position Converter is the most frequently used tool in the Mutalyzer suite.

The usage of the three latest human reference genomes is shown in Figure 7.

**Figure 7:**
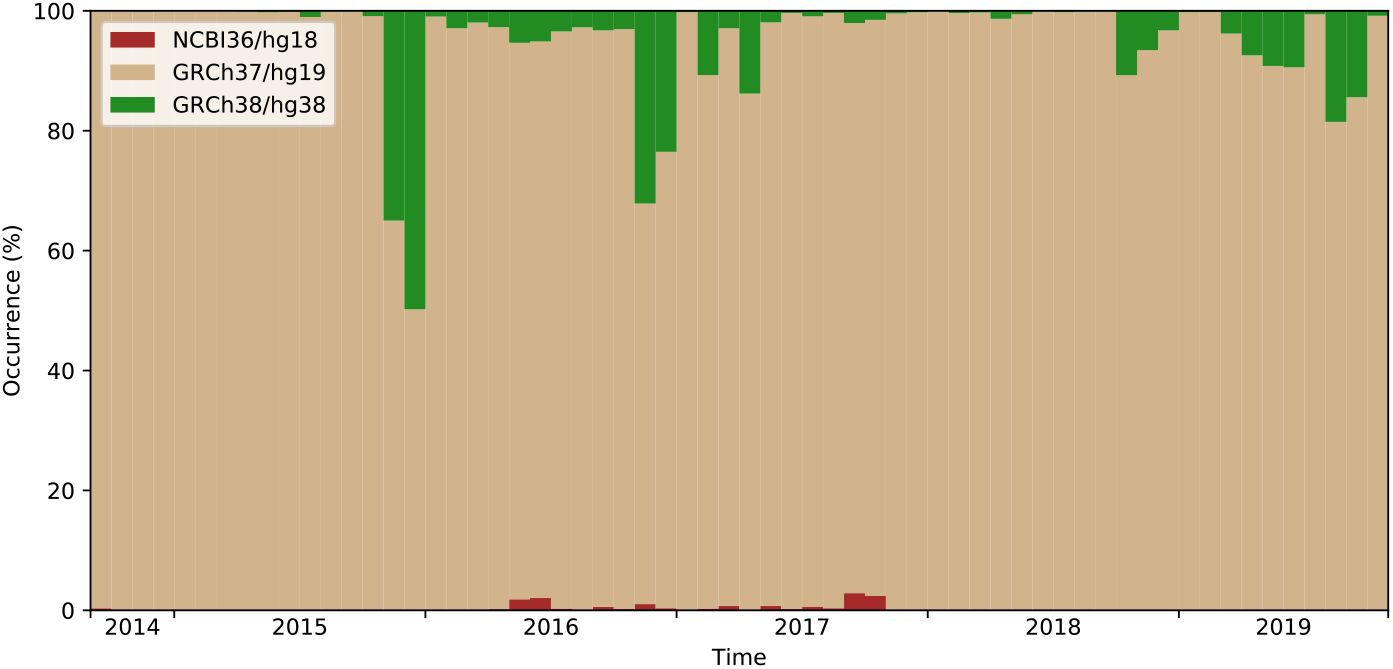
Reference genomes used in Position Converter descriptions.

Of the latest three human reference genomes, GRCh37 is the one most frequently used (94%). Although GRCh38 was released in December 2013, it is being used in only 5% of the cases. It is noteworthy that NCBI36 was last used in 2017.

## 7 Discussion

In this section we discuss common misconceptions and issues that arise from using standard reference sequence files.

### 7.1 Equivalent Descriptions

The NCBI provides reference sequence files for various genomic features at different levels [16]. Chromosomal assembly reference files (identified by the accession number prefix `NC_`) contain both the entire chromosome sequence and annotation of its corresponding features. We refer to a transcript annotated on a chromosome as *chromosomal transcripts*. Transcript reference files, or *RefSeq transcripts*, (identified by the accession number prefix `NM_` or `NR_`) are available from the same source. These references only contain the sequence and features for one transcript, they therefore do not contain any intronic sequences.

The HGVS `c.` and `n.` positioning systems make it possible to describe a variant in the context of a transcript, while using a genomic (chromosomal) reference sequence. The same notation can be used to describe a variant using a RefSeq transcript, which results in a very similar looking, but potentially very different variant description. This seemingly equivalent way of describing variants is the source of many erroneous descriptions.

Tools like the Position Converter and the Variant Validator [10], provide a way to convert between chromosomal and RefSeq transcripts descriptions. However, because RefSeq transcripts lack intronic sequences and, moreover, exonic sequences may differ between chromosomal and RefSeq transcripts, the results of such a conversion should be interpreted with great care.

A summary of the differences between chromosomal and RefSeq transcripts for in human genome builds GRCh37 and GRCh38 is shown in Table 2. It is important to note that 6.79% of the transcripts in GRCh37 and 2.52% of the transcripts in GRCh38 differ in terms of sequence content from those found in RefSeq. Moreover, in over 50% of the cases, these differences are found in the protein coding region. This is why it is important not to confuse a chromosomal transcript like `NC_123.4(NM_567.8)` with `NM_567.8`, as they may differ quite substantially.

**Table 2:** Sequence differences between chromosomal and RefSeq transcripts.

| Assembly | Total NMs<br>(#) | Different NMs<br>(#) | (%) | SNPs<br>(%) | Alleles<br>(%) | In CDS<br>(%) |
| --- | --- | --- | --- | --- | --- | --- |
| GRCh37 | 34641 | 2353 | 6.79 | 78.20 | 21.80 | 64.56 |
| GRCh38 | 56104 | 1415 | 2.52 | 82.05 | 17.95 | 52.65 |

\* Retrieved between February and April 2020.

The majority of the differences (about 80%) found in the survey described above, consist of single nucleotide variants, but there are cases in which deletions or insertions occur. These differences change the mapping of positions between a chromosomal and a RefSeq transcript sequence. Such an example is shown in Figure 8, where in the chromosomal transcript a sequence of five nucleotides is inserted between `c.131` and `c.132`. This results in a discrepancy for all positions downstream of this variant. In addition, the reading frame is changed, which leads to various consequences at the protein level.

**Figure 8:**
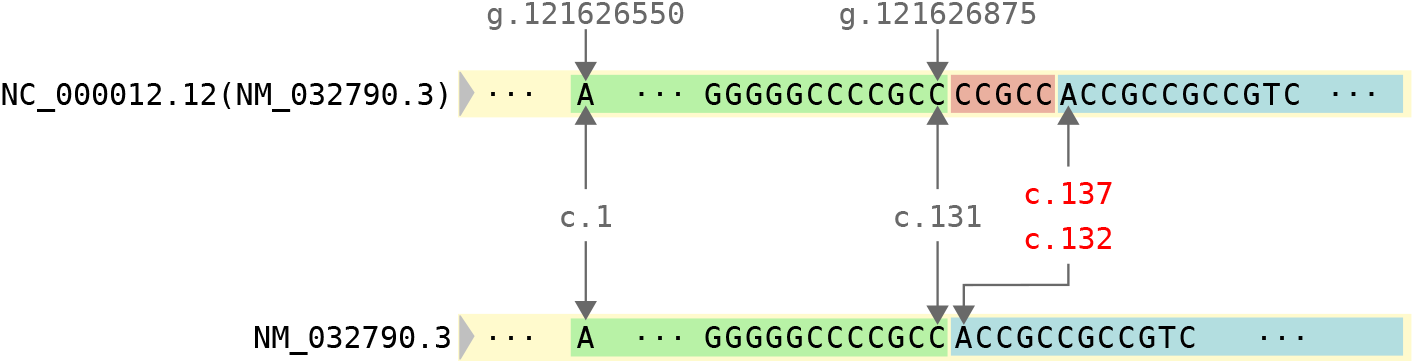
Example of different sequences between an NC and an NM.

For consistent variant dissemination we recommend using either chromosomal transcripts or RefSeq transcripts, whichever was used in the primary data analysis. We strongly advise against conversion between the two. When a transcript-oriented description is desired for variants found in Next Generation Sequencing (NGS) experiments, our advice is to use the notation `NC_123.4(NM_567.8)` instead.

### 7.2 Unstable Annotations

Mutalyzer is a fully deterministic system, which means that a given input will always yield the same result. It is possible however, that the input changes without the end user noticing, giving rise to unexpected results. A change in the reference model, like the addition or removal of features and changes in feature positions, is the most common source of such problems.

For example, version `08-APR-2018` of RefSeq file `NM_012115.3` has an exon missing compared to the one dated `11-AUG-2018` and version `06-JUN-2016` of `NC_000002.12` contains annotation for `NM_032506.2`, while the version dated `26-MAR-2018` does not. Approximately 1,000 mRNA features were removed between the two mentioned dates for `NC_000002.12` alone.

According to [17], the NCBI updates the reference identifier version number only when there is a change in the sequence. When alterations within the feature annotation section occur, there is no version update. From the presented examples, it is clear that Mutalyzer, or anyone for that matter, is not able to ensure consistent output unless the reference sequence providers maintain proper reference sequence versioning, both at the sequence and annotation level.

## 8 Support

Mutalyzer is an Open Source project available on GitHub^6^ under the GNU Affero General Public License. The GitHub issue tracker system is used for feature requests and error reporting.

The Mutalyzer mailing list^7^ is a general forum for the Mutalyzer tool suite. Messages can be posted through the Google Groups interface or by sending an email^8^. Additionally, there is a low volume mailing list where updates to Mutalyzer are announced^9^. Private questions or security related issues can be communicated via a private email address^10^.

Support on locally installed instances of the Mutalyzer tool suite, for example for the analysis of private/confidential variants, can be arranged through PhenoSystems S.A.^11^.

## 9 Conclusions and Further Research

In this paper we presented Mutalyzer 2, a tool suite created to assist geneticists in applying the HGVS variant nomenclature guidelines for consistent dissemination in clinical research, databases and scientific literature since its launch in August 2010, Mutalyzer proved its utility by processing over 133 million variant descriptions. Log statistics indicate that approximately 50% of the input descriptions were correct, while for 41% of them, either a syntactic or a semantic error was identified. In approximately 7% of the cases, Mutalyzer provided a corrected description.

For upcoming versions of Mutalyzer, support for reference sequence providers such as EMBL-EBIs Ensembl should be added. Support for descriptions that use multiple reference sequences, e.g., `g.123_124insLRG_199:g.2233_2361` is also highly desirable. Finally, the Description Extractor should be used as a central component of the Name Checker, as this will allow for disambiguation of complex allele descriptions.

## Acknowledgements

The authors thank the following people for their contributions to Mutalyzer 2 (in alphabetical order): Ivo F.A.C. Fokkema, Mark Kroon, Gerard C.P. Schaafsma and Gerben R. Stouten. This work was partly funded by the Netherlands Bioinformatics Centre (NBIC), which is supported by the Netherlands Genomics Initiative (NGI). This publication was supported by the Dutch national program COMMIT.

## Footnotes

1 http://varnomen.hgvs.org

2 “Human Mutation” and “European Journal of Human Genetics” require that compliance with HGVS nomenclature must be verified using tools such as Mutalyzer.

3 https://varnomen.hgvs.org/bg-material/consultation/svd-wg006/

4 https://mutalyzer.nl/

5 https://mutalyzer.nl/webservices

6 https://github.com/mutalyzer/mutalyzer

7 https://groups.google.com/forum/#!forum/mutalyzer

8

9 https://groups.google.com/forum/#!forum/mutalyzer-announce

10

11 http://www.phenosystems.com/www/index.php/products/mutalyzer

